## Supplementary Figures and Table for "SIRT2-Mediated ACSS2 K271 Deacetylation Suppresses Lipogenesis Under Nutrient Stress"

### Supplementary Table and Figures

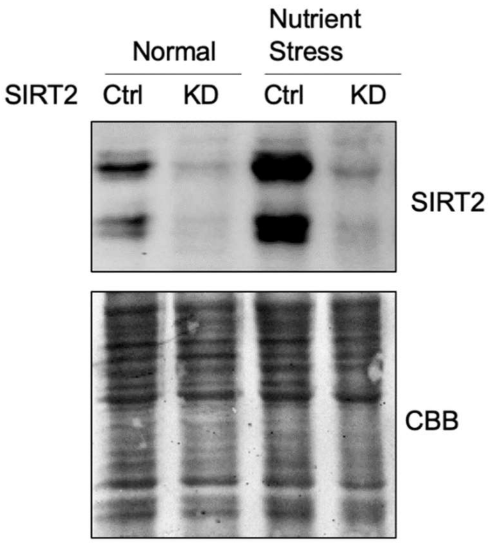


**Figure S1.** Western blot showing nutrient exhaustion increases endogenous level of SIRT2

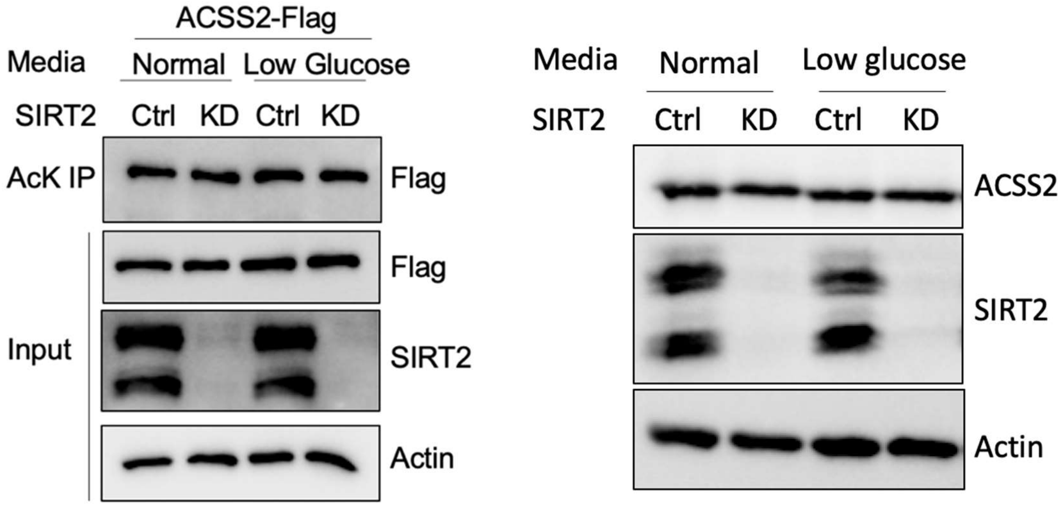


**Figure S2.** Western blot showing low glucose has no effect on ACSS2 acetylation or endogenous level of ACSS2.

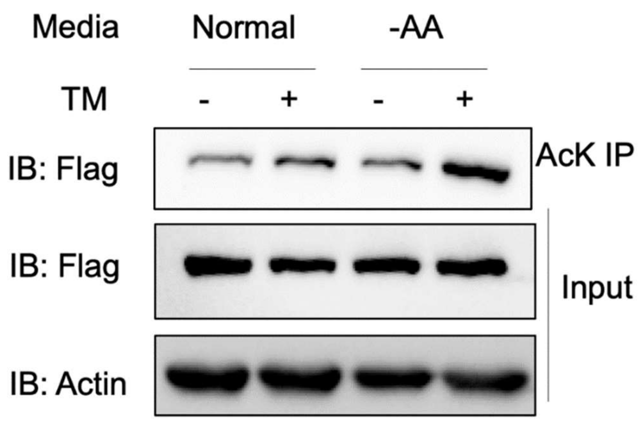


**Figure S3.** Inhibition of SIRT2 increases ACSS2 acetylation under amino acid deprivation. HEK 293T cells were treated with and without TM (SIRT2 inhibitor) and grown in normal or EBSS media. Acetylation of ACSS2 was analyzed by acetyl lysine IP and western blot for Flag.

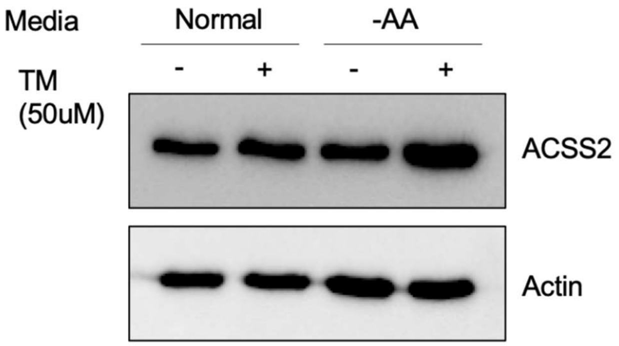


**Figure S4.** Inhibition of SIRT2 using TM results in an increase in endogenous ACSS2 in amino acid deprived media but not normal media.

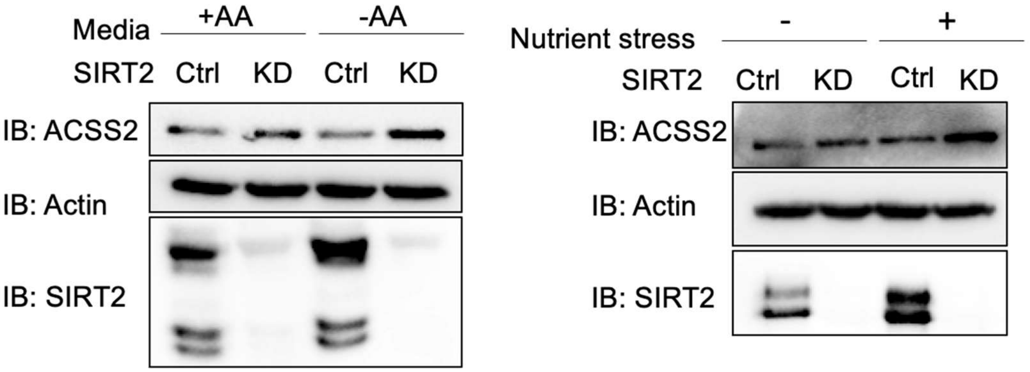


**Figure S5.** Endogenous ACSS2 levels increase when SIRT2 is knocked down under nutrient and amino acid stress. SIRT2 control and knock down A549 cells were maintained in EBSS or normal and nutrient deprived media and endogenous ACSS2 protein levels were determined by western blot.

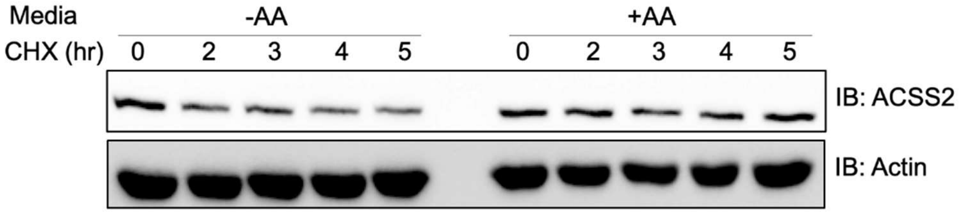


**Figure S6.** ACSS2 is destabilized in amino acid deprived media. HEK293T cells were maintained in normal and EBSS media and treated with cycloheximide (CHX) and for various time points. Endogenous ACSS2 protein levels were analyzed by western blot.

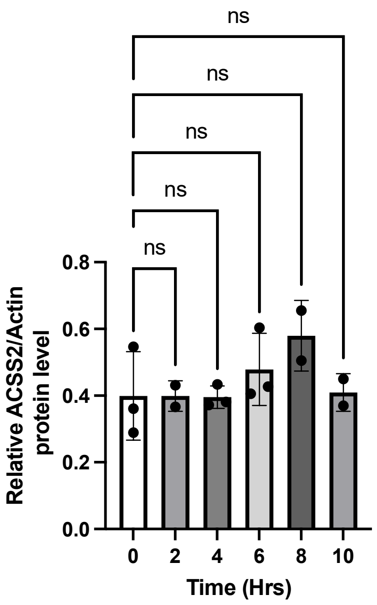


**Figure S7**. Graph of ACSS2 level for each time point for SIRT2 knockdown samples (Data from Figure 2C). There is no significant changes in ACSS2 levels.

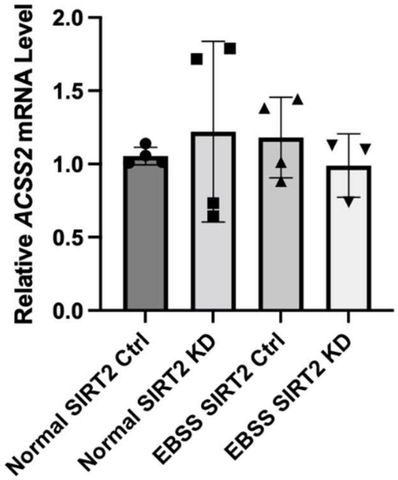


**Figure S8.** qPCR showing ACSS2 transcript levels do not change with SIRT2 knockdown or amino acid deprivation. Each data point is a biological replicate.

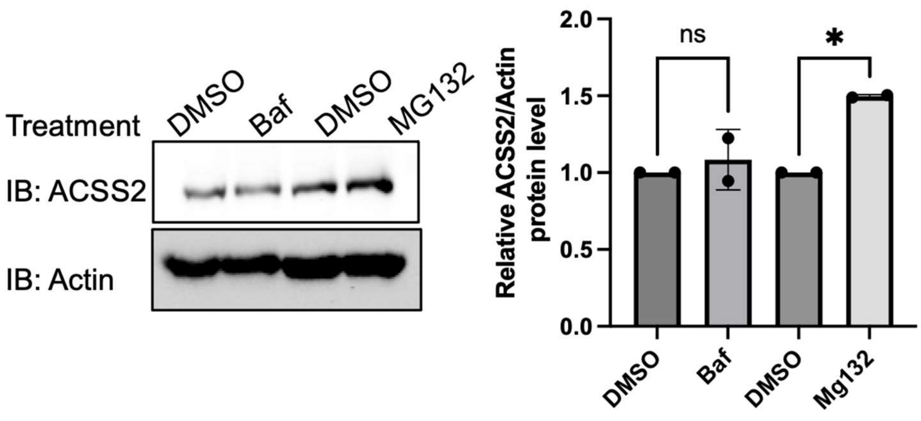


**Figure S9.** ACSS2 is degraded by the proteasome pathway. Each data point is a biological replicate.

**Table S1.** Sites on ACSS2 identified by mass spectrometry to change with SIRT2 knockdown.

| Annotated Sequence | Modifications | Modifications in Master Proteins | Abundance Ratio: (SIRT2 Knockdown) / (SIRT2 Control) | Abundances (Grouped): SIRT2 Knockdown | Abundances (Grouped): SIRT2 Control |
|---|---|---|---|---|---|
| [R].KIAQNDHDLGDMSTVADPSVISHLFSHR.[C] | 1xAcetyl [K1]; 1xOxidation [M12] | Q9NR19 1xAcetyl [K669] | | | |
| [R].LLMKFGDEPVTK.[H] | 1xAcetyl [K4]; 1xOxidation [M3] | Q9NR19 1xAcetyl [K418] | 0.959 | 97.9 | 102.1 |
| [R].AELGMGDSTSQSPPIKR.[S] | 1xAcetyl [K16] | Q9NR19 1xAcetyl [K271] | 2.433 | 141.7 | 58.3 |

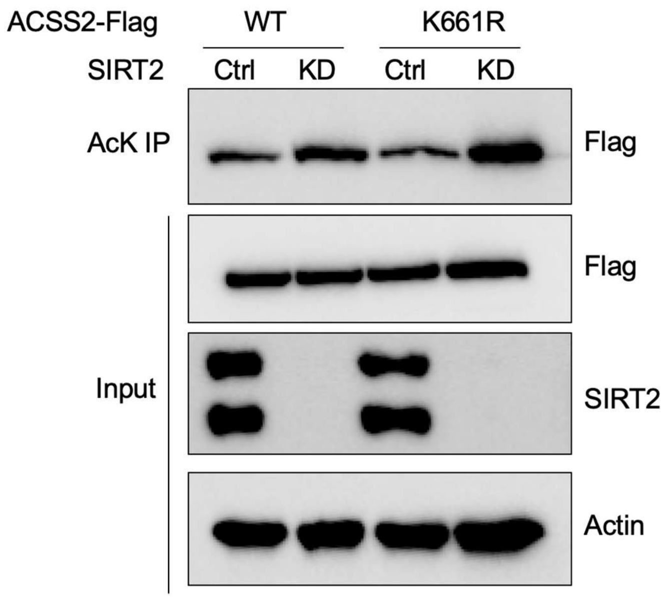


**Figure S10**. Western blot showing SIRT2 does not deacetylate ACSS2 at K661.

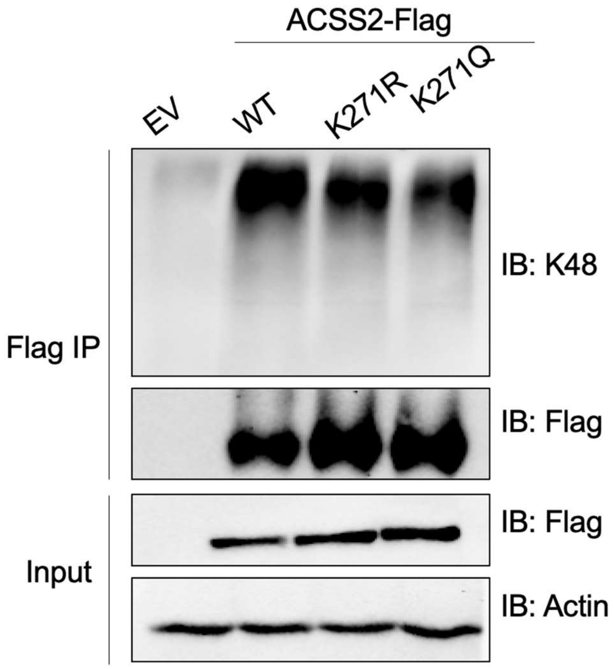


**Figure S11.** Mutation of K271 decreases ACSS2 ubiquitylation.

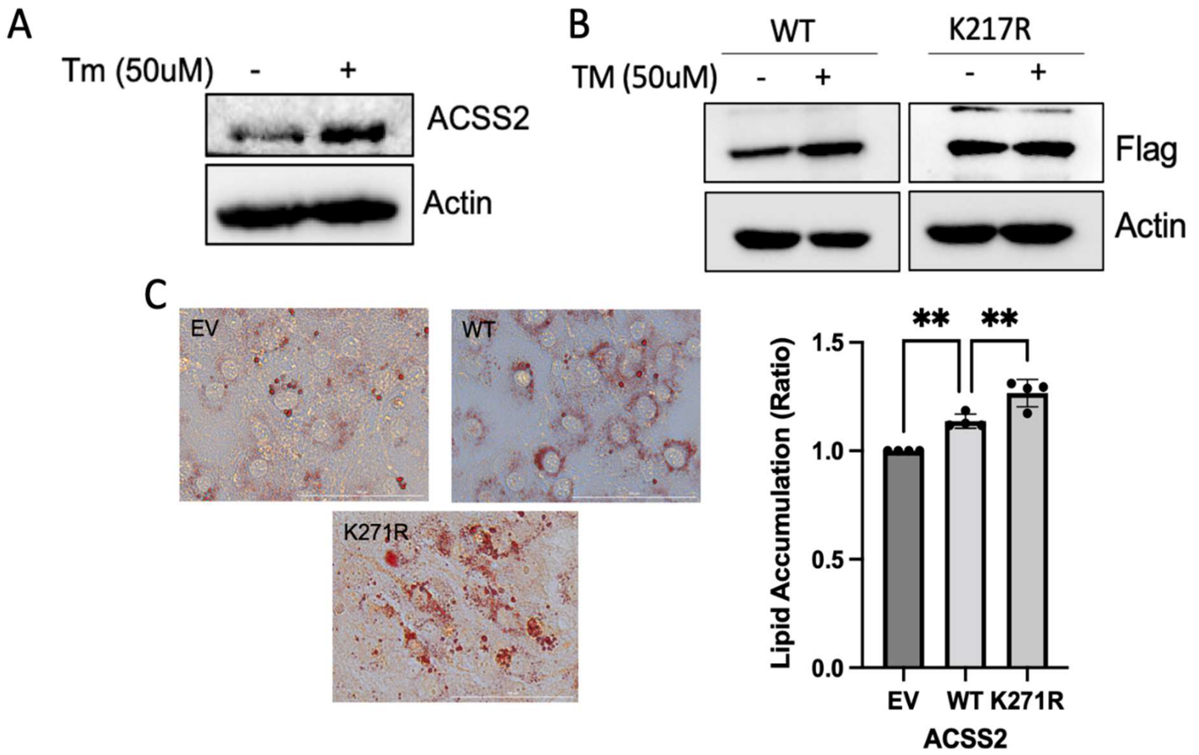


**Figure S12. Acetylation at K271 promotes increased lipid accumulation.**
(A) TM treatment in 3T3-L1 cells results in increased levels of endogenous ACSS2.
(B) Inhibition of SIRT2 with TM leads to an increase in WT ACSS2 levels but does not affect the levels of the K271R mutant.
(C) Oil Red O quantification shows that cells expressing the K271R mutant accumulate higher levels of lipids.

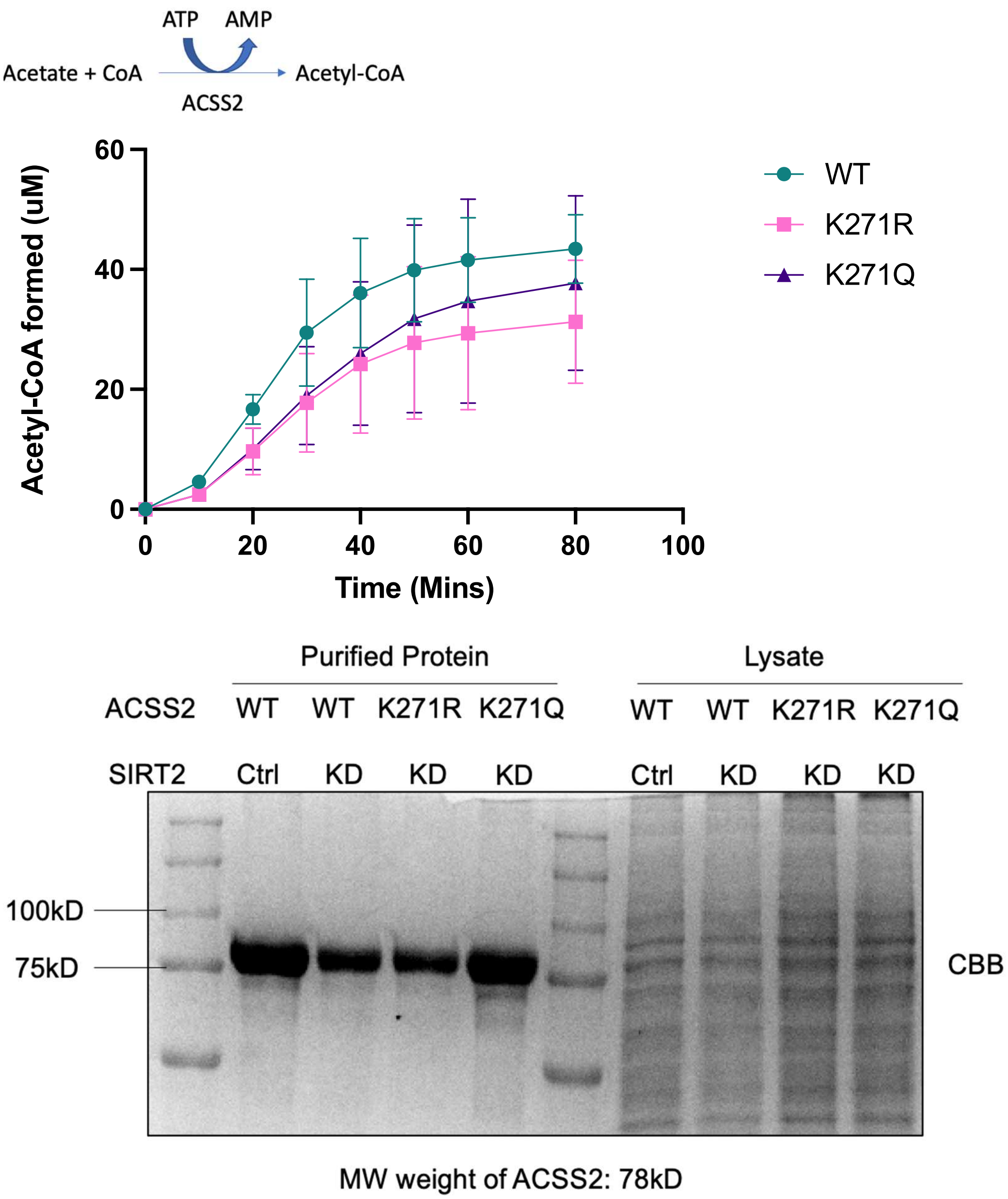


**Figure S13**. ACSS2 WT and mutant activity assay. Overexpressed ACSS2-Flag WT, K271R and K271Q was purified from SIRT2 knockdown HEK293T cells. ACSS2 activity was determined by measuring the formation of acetyl-CoA from acetate and CoA over time.

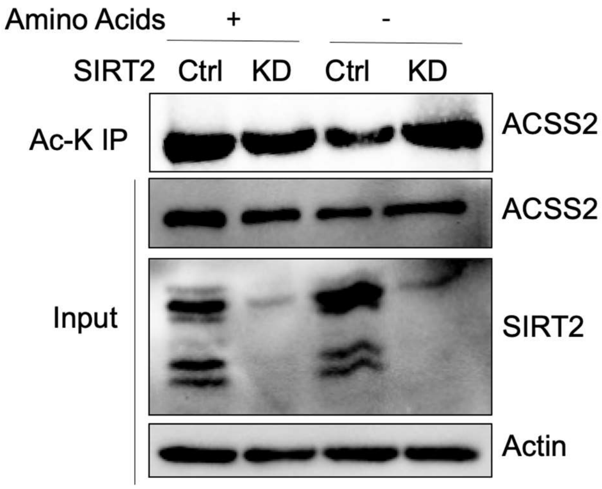


**Figure S14**. ACSS2 acetylation decreases under amino acid deprivation.

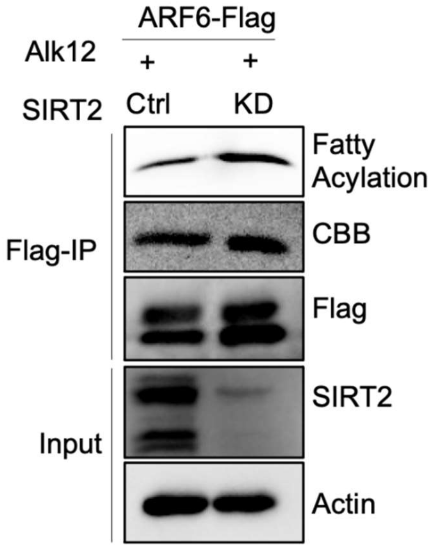


**Figure S15**. SIRT2 removes ARF6 myristylation. Alk12 labeling results of overexpressed ARF6 in SIRT2 control and knockdown cells showing that SIRT2 demyristylates ARF6 in HEK293T cells as indicated by the increased myristylation signal with SIRT2 knockdown.

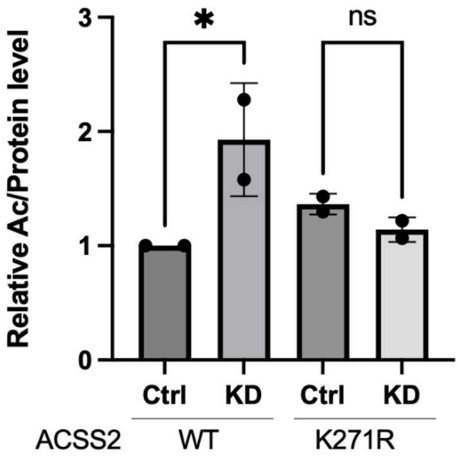


**Figure S16**. Quantification of WT and K271R ACSS2 in SIRT2 control and knockdown cells.
